# The pitfalls of using Gaussian Process Regression for normative modeling

**DOI:** 10.1101/2021.05.11.443565

**Authors:** Bohan Xu, Rayus Kuplicki, Sandip Sen, Martin P. Paulus

## Abstract

Normative modeling, a group of methods used to quantify an individual’s deviation from some expected trajectory relative to observed variability around that trajectory, has been used to characterize subject heterogeneity. Gaussian Processes Regression includes an estimate of variable uncertainty across the input domain, which at face value makes it an attractive method to normalize the cohort heterogeneity where the deviation between predicted value and true observation is divided by the derived uncertainty directly from Gaussian Processes Regression. However, we show that the uncertainty directly from Gaussian Processes Regression is irrelevant to the cohort heterogeneity in general.

## Introduction

In case-control studies, participants are assigned labels and classified into one or more categories based on their similarities or common criteria, with little consideration for the heterogeneity within each cohort. Meanwhile, normative modeling is becoming increasingly popular. In a normative model, each observation is quantified as a normalized deviation with respect to the cohort heterogeneity. The growth chart [1, 2] is an example normative model as shown in Fig. 1, where a series of percentile curves (normalized deviation) illustrate the distribution of selected body measurements in children. Another widely-used measure for normalized deviation is the *z*-score, which is calculated by dividing the difference between an observation and the reference model, i.e., residual, by a standard deviation that represents local heterogeneity and assumes residuals are Gaussian distributed locally.

**Fig 1.**
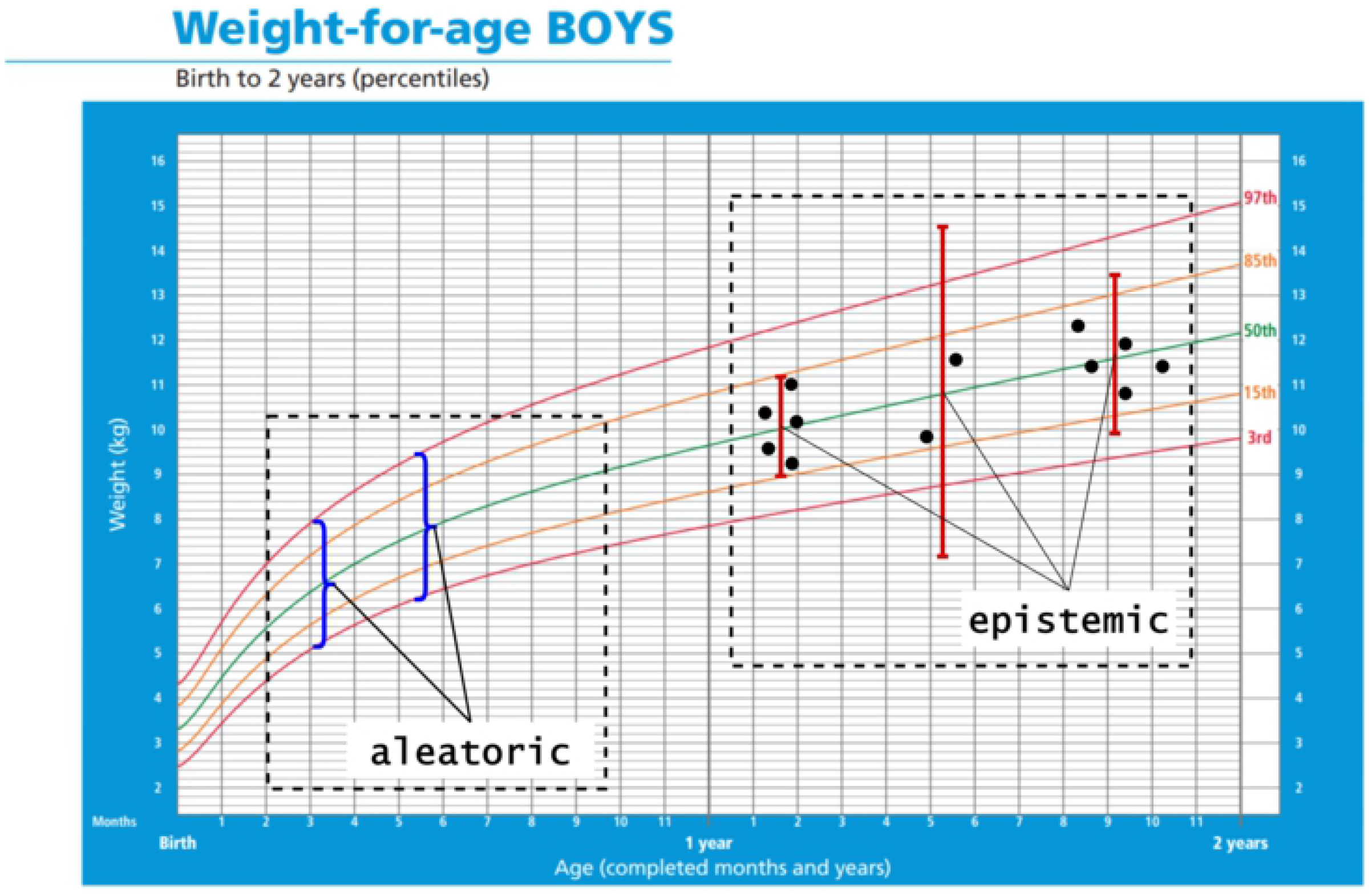
Weight-for-Age Boys: Birth to 2 years [3]. The percentiles show the distribution of weights in boys form birth to 2 years. Black dots: observations; red error bars: epistemic uncertainty; blue curly brackets: aleatoric uncertainty.

The uncertainty sometimes can be classified into two categories: epistemic and aleatoric uncertainties. Epistemic uncertainty is known as systematic uncertainty and is due to things one could in principle know but do not in practice; aleatoric uncertainty is known as statistical uncertainty and is representative of unknowns that differ each time we run the same experiment [4]. Epistemic uncertainty is often introduced by the limited dataset size and can be reduced by adding more observations. On the other hand, aleatoric uncertainty represents a character of heterogeneity in the underlying distribution itself which is unrelated to sample size, so it cannot be reduced by modifying the dataset, and this is the heterogeneity a normative model should measure. As shown in Fig. 1, larger number and density of data points (black dots) reduce the epistemic uncertainty (red error bars), while the aleatoric uncertainty (blue curly brackets) is unrelated to the sample size or distribution. The confidence intervals obtained from most statistical tests and advanced machine learning models only capture epistemic uncertainty, while a normative model is designed to capture the aleatoric uncertainty.

Gaussian Process Regression (GPR) has been widely used in many domains. Schulz et al. [5] presented a tutorial on the GPR with the mathematics behind the model as well as several applications to real-life datasets/problems. Tonner et al. [6] developed a GPR based model and testing framework to capture the microbial population growth and shown their proposed approach outperformed primary growth models. Banerjee et al. [7] and Raissi et al. [8] introduced two novel approaches to improve the efficiency of GPR in “big data” problems.

However, some previous research implemented the GPR as a normative modeling approach and utilized the derived prediction variance to model the cohort heterogeneity. Ziegler et al. [9] attempted to build a normative model for diagnosing mild cognitive impairment and Alzheimer’s disease based on the normalized deviation of predicted brain volume from GPR. Marquand et al. [10] used delay discounting as covariates and reward-related brain activity derived from task Functional Magnetic Resonance Imaging (fMRI) as the target variable with GPR and extreme value statistics to identify the participants with Attention-Deficit/Hyperactivity Disorder (ADHD). Wolfers et al. [11] investigated the deviation of brain volume in an ADHD cohort from healthy control group (HC) with respect to age and gender, and they also explored the heterogeneous phenotype of brain volume for schizophrenia and bipolar disorder with GPR [12]. Zabihi et al. [13] studied Autism Spectrum Disorder (ASD) regarding the deviation of cortical thickness via a similar methodology.

In this paper, we introduce some background knowledge related to GPR. We then present a rigorous mathematical derivation and several examples to demonstrate that the variance from GPR cannot be used in a normative model alone. In the last section, we discuss the difficulties and disadvantages of modeling the cohort heterogeneity by modifying original GPR variance, and a misunderstanding existed in previous research.

## Materials and methods

### Gaussian Process Regression

The relation between the observation and the predictive model usually can be expressed as

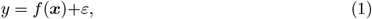

where *y* is the observation (output), *f* (·) represents the predictive model, ***x*** is a vector of independent variables (input) corresponding to the output *y*, and *ε* is the noise term which follows a normal distribution 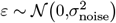. Gaussian Process Regression (GPR) assumes a zero-mean normal distribution over the predictive model

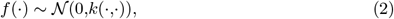

where *k*(·,·) is some covariance (kernel) function. Given the training set input ***X*** and testing set input ***X***_∗_, since both of them follow the same distribution, we have

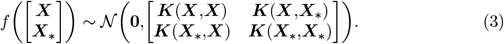

According to the Eq. 1, the observation follows the summation of these two normal distributions

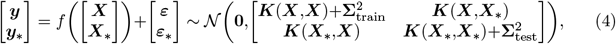

where 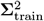 and 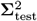 are two diagonal matrices that represent the variance of observation noise in training and testing sets, and the diagonal elements equal 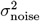. By the rules of conditional Gaussian distribution, the prediction of testing set ***y***_∗_ follows a normal distribution 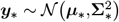, where ***µ***_∗_ and 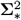 are defined as [14, 15]

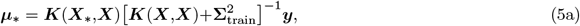

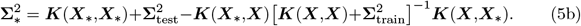

### Kernel Trick

Similar to Support Vector Machines (SVM), the kernel trick can also be implemented with GPR to project the input of data from the original space into a same or higher dimensional feature space via some mapping function *z*(·). Given a pair of inputs (***x***_1_,***x***_2_), the kernel function calculates the inner product of the coordinates in the feature space, i.e., *k*(***x***_1_,***x***_2_) = ***z***(***x***_1_)***z***(***x***_2_)^*T*^ [16, 17]. The kernel trick avoids the expensive computation of calculating the coordinate in the feature space for each input. We use the linear kernel and Radial Basis Function kernel (RBF) as examples to illustrate this advantage.

### Linear Kernel

The linear kernel is non-stationary and the simplest kernel, which is defined as

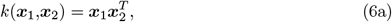

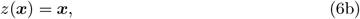

where the input is projected into a feature space according to Eq. 6b, and the feature space is the original space.

### Radial Basis Function Kernel

The RBF kernel is a stationary kernel, which is also widely used and defined as [17]

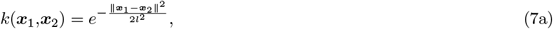

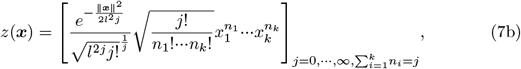

where *l* is a free scaling parameter. The RBF kernel projects the input from the original space onto an infinite dimensional feature space where the mapping is defined by Eq. 7b. It is impossible to exactly compute the coordinates in an infinite dimensional space, while Eq. 7a still allows straightforward computation of the inner product for coordinate pairs in that feature space.

### Estimated Uncertainty for GPR

One benefit of using GPR to build a data-driven model is the predictions are associated with the derived variances as shown in Eq. 5. However, we need to emphasize that this variance is only related to the kernel function *k*(·,·) and distribution/coordinate of training set input ***X***, i.e., it cannot be utilized in a normative model approach alone to capture the variance introduced by the conditional distribution *V ar*(*y*|***x***).

We better illustrate and verify this statement through simplifying the Eq. 5b. Since any kernel function *k*(·,·) can be written as the inner product of a coordinate pair in the feature space by some mapping function *z*(·), we present our derivation in a general format. We define a variable ***x***_∗_ which represents a testing input, then Eq. 5b can be written as^1^

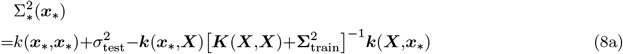

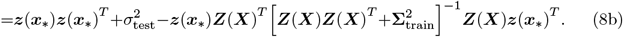

Applying Singular Value Decomposition (SVD) on 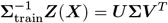, Eq. 8 is reformulated as^2^

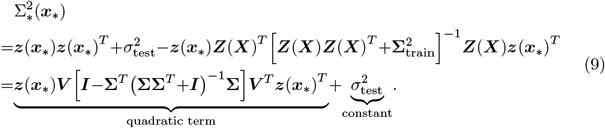

After simplification, the variance is reformulated as Eq. 9, which is a summation of a quadratic term for ***z***(***x***_∗_) and a constant represents the noise, **Σ** and ***V*** are constant matrices where the values are fully depended on training input ***X***, training noise **Σ**_train_, and mapping function *z*(·) or kernel function *k*(·,·).^3^

### Modification of Uncertainty from GPR

Regarding Eq. 9, the variance calculated via Eq. 5b is purely depended on kernel function and training data input, thus it is only able to capture the epistemic uncertainty which could be reduced by modifying or adding training data. The derived variance from GPR could be extended to model the heterogeneity *V ar*(*y*|***x***) for a normative model by adding an aleatoric variance term into Eq. 5b

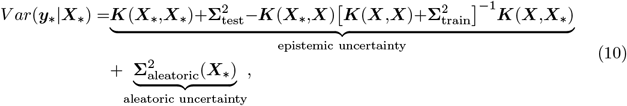

where 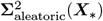 represents the data character of heterogeneity in output at given locations on the input space. This formula, however, is not implemented in any previous research as we know and we will discuss the difficulties and disadvantages in estimating the aleatoric uncertainty later.

## Results

We apply the unmodified GPR (Eq. 5) on several synthetic datasets where both input *x* and output *y* are one dimensional to facilitate visualization. Although the presented results are based on one dimensional input *x*, they are generalizable to any dimensional input. The selected kernels are linear and RBF kernels, and we present the results of two scenarios with known and unknown noise levels.^4^

### Dataset

Four synthetic datasets are generated and plotted in Fig. 2, and each of them contains 1000 points with a noise level of σ_noise_ = 0.05. Four other undersampled datasets are plotted in Fig. 3, each of which contains 1000*×*5% = 50 points.

**Fig 2.**
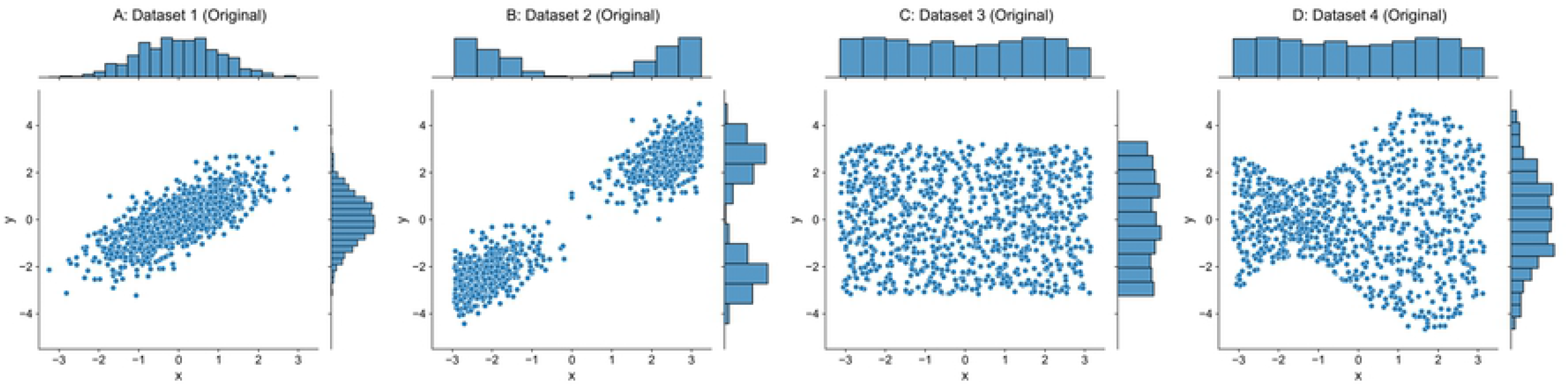
Original Datasets.

**Fig 3.**
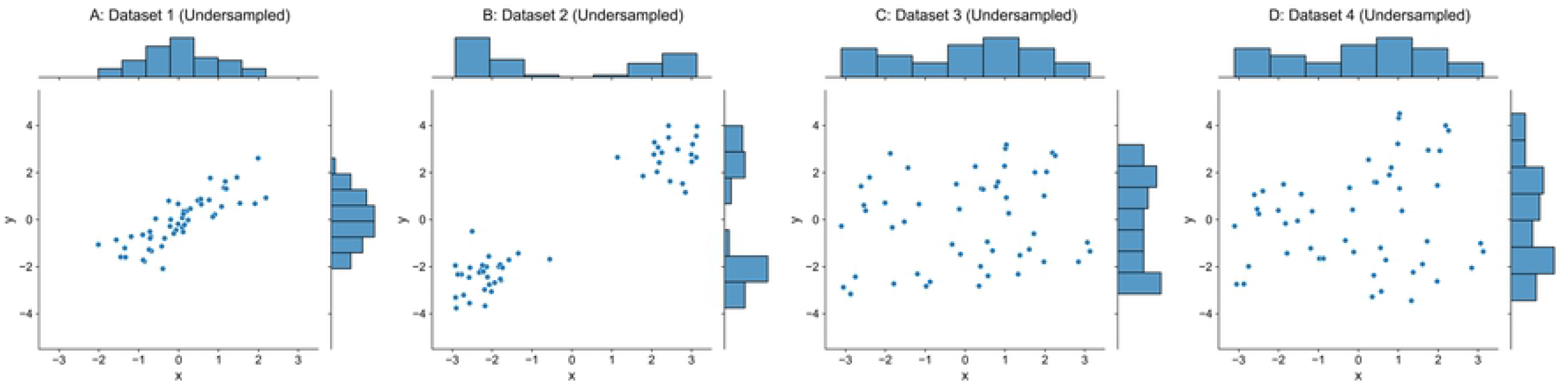
Undersampled Datasets.

In Dataset 1, both input ***X*** and output ***y*** follow a Gaussian distribution *N* (0,1^2^) and are correlated with a Pearson coefficient of 0.75. Dataset 2 is transformed from Dataset 1, which moves the set of points where *x* ≥ 0 in Dataset 1 along the line *y* = *x* until the maximum input in that set equals 0, and moves the remaining points where *x <* 0 in Dataset 1 until the minimum input is 0. Dataset 3 has input ***X*** and output ***y*** uniformly distributed over a half-open interval [−*π,π*). Output ***y*** of Dataset 4 is obtained by multiplying a factor function over output ***y*** from Dataset 3, which is defined as *f* (***x***) = sin(***x***)*/*2+1 and ***x*** is the corresponding input. We should note that the inputs ***X*** of original Datasets 3-4 are exactly same as shown in Fig. 2C-2D, and the inputs ***X*** of corresponding undersampled Datasets 3-4 are also identical as shown in Fig. 3C-3D.

### GPR with Known Noise Level

#### Linear Kernel

The regression surface of GPR with linear kernel is a hyperplane and the variance is a quadric hypersurface defined by Eq. 9 in feature/original spaces, where the hyperplane always passes the origin, the variance is a function only with respect to the coordinate of testing input ***x***_∗_ and a unique minimum is located at ***x***_∗_ = **0**. Figs. 4-5 present results for GPR with linear kernel on the one dimensional synthetic datasets, where top sub-figures plot the reference models/predictions (red lines) overlapped on the data (blue dots), middle sub-figures show the derived variances (blue curves) across the original input space, and the bottom sub-figures shows the corresponding “*z*-score” for training set which is computed via Eq. 11 if the residual (*y*−*y*_reference_) is mistakenly normalized by standard deviation Σ directly from GPR (Eq. 5b).

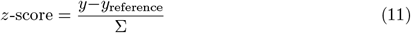

**Fig 4.**
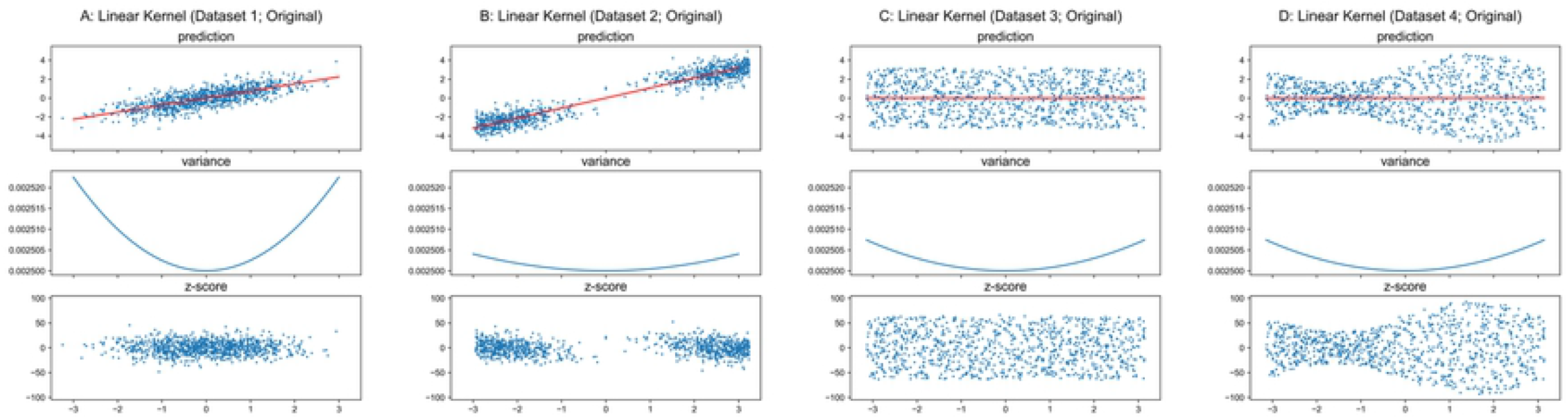
GPR with Linear Kernel on Original Datasets.

The mapping function of linear kernel projects an input to itself (Eq. 6b), and for one dimensional input, Eq. 9 can be further reduced to^5^

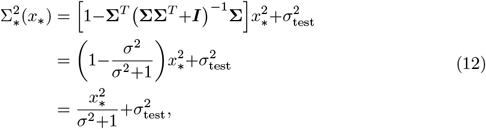

where σ is a scalar and equal to the only one singular value of 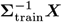. As shown in Figs. 4-5, the variance is a univariate function of coordinate of the testing input *x*_∗_ where the shape is a quadratic curve, and the global minimum is always located at *x*_∗_ = 0 with a value of 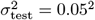 as Eq. 12 formulated. The result of GPR with linear kernel presents a good example which illustrates the derived variance from GPR does not model the conditional variance *V ar*(*y*|***x***), thus corresponding *z*-score cannot be utilized as a normalized deviation in a normative model.

As previously mentioned, the predicted variance for testing set from GPR only depends on the training set input and the kernel function. As the original as well as the undersampled Datasets 3-4 have identical inputs ***X***, the variance curves in Figs. 4C-4D and Figs. 5C-5D are respectively identical.

**Fig 5.**
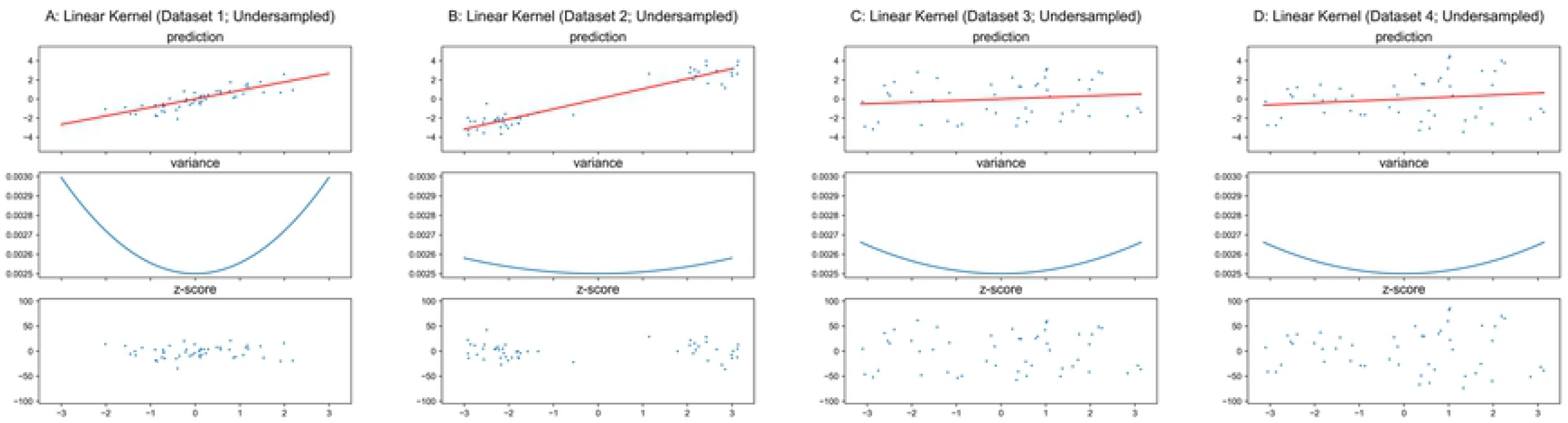
GPR with Linear Kernel on Undersampled Datasets.

#### RBF Kernel

Unlike the linear kernel, RBF kernel mapping function (Eq. 7b) defines a feature space which is different from the original space. Therefore, regarding the original space, the regression surface is no longer a hyperplane and the variance is no more a quadric hypersurface for the RBF kernel.^6^ Because the mapping function of RBF kernel is very complicated, we only briefly describe the characteristics of the regression surface and variance in the original space. For a test input ***x***_∗_, the prediction is a summation of discounted outputs of all training points where each corresponding discount factor is determined by the Euclidean distance between ***x***_∗_ and that training input, and the predicted value converges to 0 if ***x***_∗_ is far away from all training inputs. On the other hand, the variance depends only on the density of training inputs at ***x***_∗_, and higher density results in lower variance. Therefore, the variance of GPR with RBF kernel is related of the relative location to the training inputs rather than the absolute location specified by coordinate.

The results for GPR with RBF kernel applied to these synthetic datasets are shown in Figs. 6-7.^7^ As shown in Figs. 6-7, the variance is unrelated to the conditional variance *V ar*(*y* | *x*). Therefore, *z*-scores based on this model do not represent normalized deviation. However, unlike the quadratic curves whose unique minimum is always located at *x* = 0 in Figs. 4-5 for linear kernel, the variance function of GPR with RBF kernel regarding the original input space is related to the distribution of training input ***X***. The denser inputs at the middle of Dataset 1 and two ends of Dataset 2 lead to lower variances at those locations in Figs. 6A-6B, while the uniformly distributed inputs of Datasets 3-4 result in relatively flat curves in Figs. 6C-6D. According to Eq. 7a and given an arbitrary input ***x***_∗_, the RBF kernel function returns a larger value for a point in ***X*** that is closer to ***x***_∗_, and ***k***(***x***_∗_, ***X***) and ***k***(***X***,***x***_∗_) have more large elements if ***x***_∗_ is close to more points in ***X***. Both result in the decrease of the value for Eq. 8a, i.e., to smaller variance.^8^

**Fig 6.**
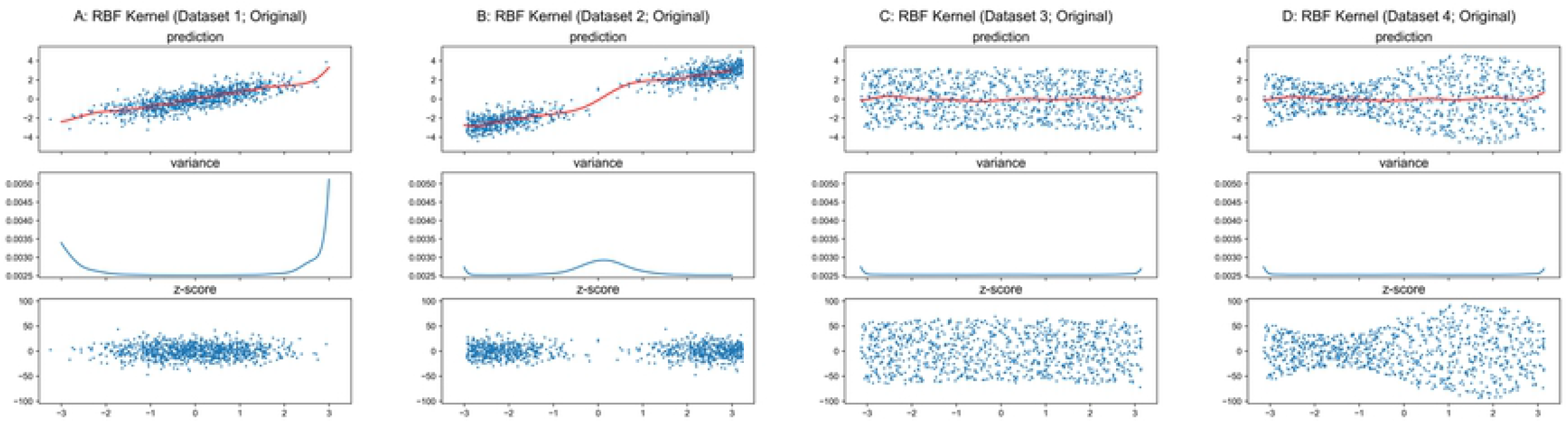
GPR with RBF Kernel on Original Datasets.

Similar to the result for the linear kernel, the theoretical minimum of variance is 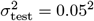,^9^ and the variance curves are exactly identical in Figs. 6C-6D and Figs. 7C-7D respectively.

**Fig 7.**
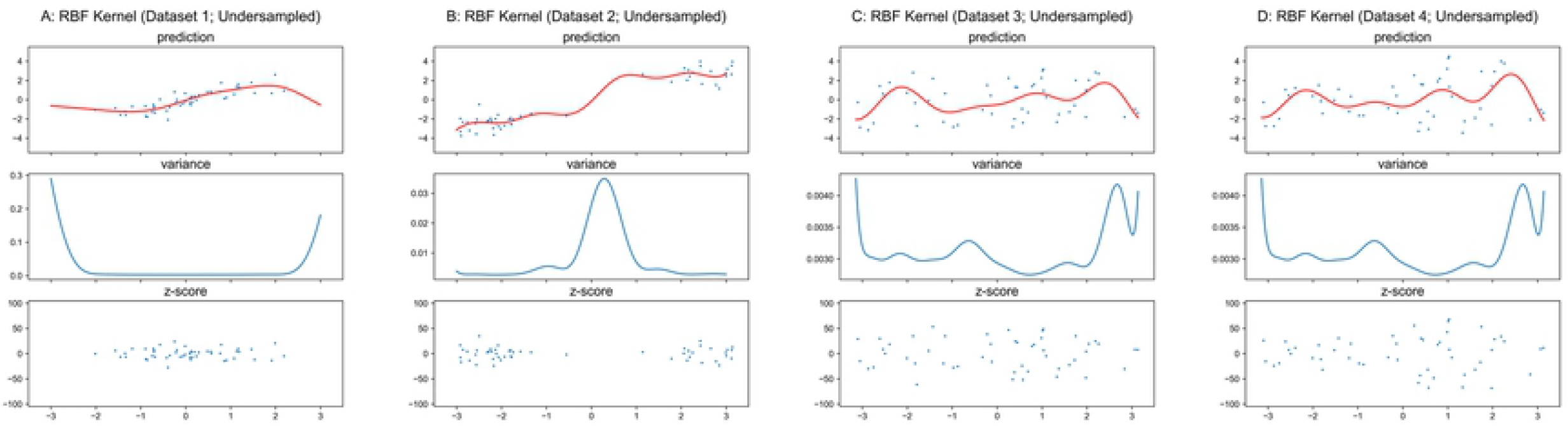
GPR with RBF Kernel on Undersampled Datasets. Scales of Y-axis for variance plots are different.

### GPR with Unknown Noise Level

The noise level can be included as a hyper-parameter when it is unknown. However, the derived variance from GPR still does not model the heterogeneity *V ar*(*y*|***x***), although it could be a good approximation in some special cases.

As the basic properties of linear and RBF kernels have been introduced, a hybrid kernel is utilized in the following analysis which is defined as

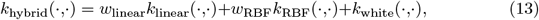

where *w*_linear_ and *w*_RBF_ represent adjustable weights on linear and RBF kernels, and *k*_white_(·,·) refers to a white-noise kernel that represents the 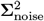.^10^ The Eq. 5b is reformulated as

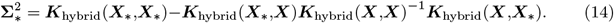

Original Datasets 3-4 in Figs. 2C-2D are prefect for testing whether a model captures the heterogeneity *V ar*(*y*|***x***), as the large number of instances and uniformly distributed data over the input space lead to negligible epistemic uncertainty in certain input range, and the true reference model *y* = 0 is very simple.^11^ The hyper-parameters are tuned by maximizing the likelihood *P* (***y***|***X***,***θ***), where ***θ*** represents all hyper-parameters in the model. The results are plotted in Fig. 8, and the optimized hyper-parameters are listed in Table 1 as well as the overall variances of residual *V ar*(*y*−*y*_reference_).

**Table 1.** Optimized Hyper-parameters for Hybrid Kernel on Original Datasets 3-4.

| | $w_{\text{linear}}$ | $w_{\text{RBF}}$ | $l$ | $\sigma_{\text{noise}}^2$ | $Var(y-y_{\text{reference}})$ |
| --- | --- | --- | --- | --- | --- |
| Dataset 3 | 7.29e-8 | 4.88e-10 | 2.94e2 | 3.29 | 3.29 |
| Dataset 4 | 1.35e-7 | 1.98e-17 | 1.54e-5 | 3.64 | 3.64 |

**Fig 8.**
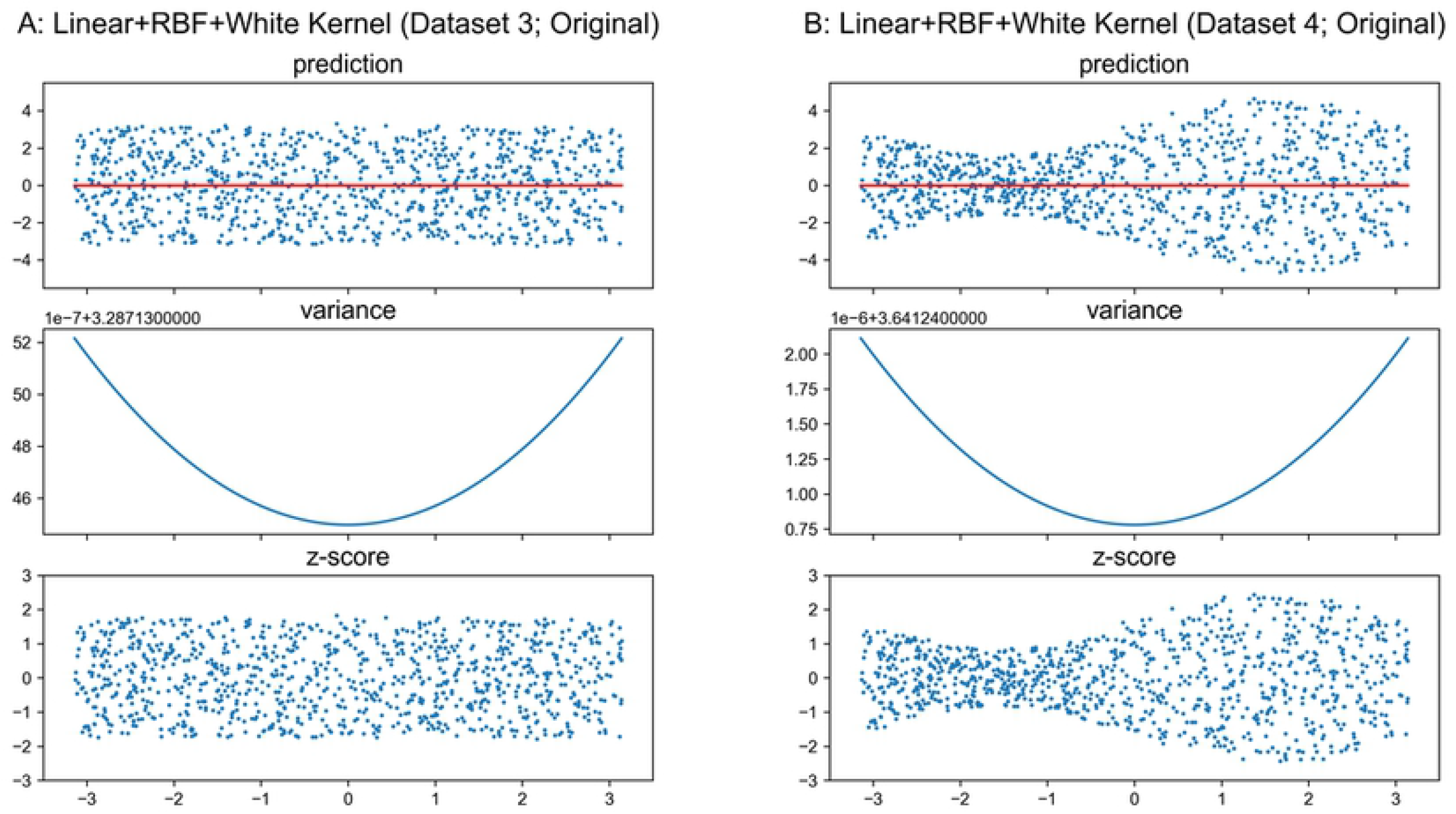
GPR with Hybrid Kernel on Original Datasets 3-4. Scales of Y-axis for variance plots are different.

As shown in Fig. 8, the GPR accurately estimates the reference models, i.e., *y*_reference_ ≈ *y*_reference,true_. The variance curves are nearly quadratic, since the *w*_linear_ is relatively larger than *w*_RBF_ while *w*_RBF_ is not exact zero as listed in Table 1. However, the domination of *k*_white_(·, ·) over *k*_linear_(·, ·) and *k*_RBF_(·, ·) due to small optimized weights flattens the curves, i.e., the value of the curve is almost constant over the plotted input range in this example. Particularly, the 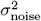 is very close to the overallresidual variance *V ar*(*y*−*y*_reference_), and the explanation will be presented later.Therefore, 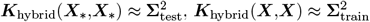, ***K***_hybrid_(***X***_*_,***X***) ≈ **0** and ***K***_hybrid_(***X***,***X***_*_) ≈ **0**, which result in 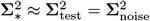.

Regarding Eq. 1, since the noise is included as a tunable hyper-parameter without any constraints, the optimizer will adjust reference model *f* (·) as well as bias 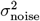 to *V ar*(*y* −*y*_reference_) to maximize the likelihood *P* (***y***|***X***,***θ***). Even the *σ*_noise_ refers to the observation noise level in GPR while the optimizer handles it as a variable without considering its meaning in a model.

In Dataset 3, 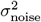 is biased to the overall residual variance *V ar*(*y*−*y*_reference_), and *V ar*(*y* − *y*_reference_) is well matched with the homoskedastic heterogeneity *V ar*(*y* |***x***). So the *z*-scores plotted in Fig. 8A show the GPR works as a normative model approach in this special case. However, in Dataset 4, 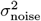 is also biased to the overall residual variance *V ar*(*y* −*y*_reference_), while *V ar*(*y* −*y*_reference_) does not approximate the heteroskedastic heterogeneity *V ar*(*y* ***x***). So the *z*-scores plotted in Fig. 8B do not represent a measure of normalized deviation in general.

## Discussion

Although GPR could be extended and to model the heterogeneity as presented in this work, it is either: (1) hard to estimate the aleatoric uncertainty accurately when the data are sparse, e.g., at the middle of Dataset 2; or (2) unnecessary to model the conditional variance by Eq. 10 when the data are dense, e.g., Datasets 3-4. One approach to estimate 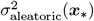 is using the sliding window technique, but it is hard to choose the window size for each dimension of input. For Scenario 1, even if the optimal window sizes can be obtained, it is hard to accurately estimate 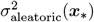 when the window centered at ***x***_*_ only covers a small number of training data points, e.g., ***x***_*_ is far away from all points in ***X***. If the window centered at ***x***_*_ covers a large number of training data points, e.g., Scenario 2, *V ar*(*y*|***x***_*_) should almost equal 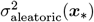 and epistemic uncertainty is insignificant. Then *V ar*(*y*|***x***_*_) can be simply approximated as a local variance over a space defined by the window.

Another misunderstanding we found in the literature is interpreting the noise term 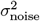 as aleatoric uncertainty. When the observation noise is considered as a hyper-parameter, it will likely bias the overall residual variance *V ar*(*y* − *y*_reference_). The overall residual variance is a good approximation of homoskedastic aleatoric uncertainty *V ar*(*y* ***x***). It is, however, not valid for cases with heteroskedastic residuals, which is the main motivation for using normative modeling. Although the value of the noise term is biased to estimate overall residual variance during the optimization, the mathematical/physical meanings are pre-defined by the model. Moreover, in homoskedastic aleatoric uncertainty cases, further investigation is needed to verify whether 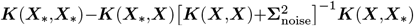 will still be a good approximation of epistemic uncertainty with such a biased estimation of observation noise level.^12^

## Conclusion

In this paper, we present the mathematical derivation with a general formula to demonstrate that the derived prediction variance from GPR does not model the heterogeneity *V ar*(*y* | ***x***), which in general is necessary for a normative model. GPR with a linear kernel and an RBF kernel are used as examples to illustrate this statement on one dimensional input datasets. Overall, the derived variance from GPR cannot be utilized in a normative model alone.

## Supporting information

**S1 Appendix. This file contains Eq. S1, Fig. S1, and Table S1**. Eq. S1, a detailed derivation for Eq. 9. Fig. S1 and Table S1, results for modified Datasets 3-4.

## Acknowledgments

We would like to express our appreciation to the anonymous reviewers for insightful discussions and feedback that have improved our study and manuscript. Also, special thanks to Laureate Institute for Brain Research and the University of Tulsa for supporting this research.

## Footnotes

1 σ_noise_ = σ_train_ = σ_test_

2 **Σ** is a diagonal matrix and the elements on the diagonal are the singular values of 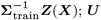; ***U*** and ***V*** are orthogonal; a detailed derivation is presented in Appendix.

3 Eq. 8b is also in the format of a summation for a quadratic term of ***z***(***x***_∗_) and a noise term.

4 These results are produced by Python 3.8.7 and scikit-learn 0.24.0; although we have not tested, it should work with other version of Python and package as well. The code for this paper is available at: https://github.com/nidaye1999/normative-model-GPR.

5 *z*(*x*) = *x* is an one dimensional vector (scalar) in this example, **Σ** is a *m×*1 matrix where Σ_1,1_ equals σ and the rest elements are 0s, ***V*** is an 1*×*1 matrix and the only one element equals 1.

6 In the feature space, regression surface is always a hyperplane and variance is always a quadric hypersurface for any kernels.

7 The value of hyper-parameter *l* in Eq. 7a does not affect the main idea of this paper, thus we used a fixed value of 1 instead of hyper-parameter optimization.

8 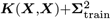 is a symmetric positive definite matrix.

9 It is a theoretical lower limit, the smallest variance for a given dataset is very likely greater than 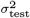.

10 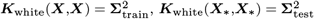,***K***_white_(***X***,***X***_*_) = **0**, and ***K***_white_(***X***_∗_, ***X***) = **0**.

11 Two datasets with quadratic reference models are presented in Appendix.

12 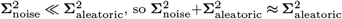 in Eq. 10.

